# Comparative performance of novel viral landscape phylogeography approaches

**DOI:** 10.1101/2025.03.24.645003

**Authors:** Simon Dellicour, Fabiana Gámbaro, Maude Jacquot, Sebastian Lequime, Guy Baele, Marius Gilbert, Oliver G. Pybus, Marc A. Suchard, Philippe Lemey

**Affiliations:** Spatial Epidemiology Lab (SpELL), Université Libre de Bruxelles, Brussels, Belgium; Department of Microbiology, Immunology and Transplantation, Rega Institute, KU Leuven, Leuven, Belgium; Interuniversity Institute of Bioinformatics in Brussels, Université Libre de Bruxelles, Vrije Universiteit Brussel, Brussels, Belgium; Ifremer, RBE-ASIM, Unité Adaptations Santé des Invertébrés Marins, Avenue de Mus de Loup, 17390 La Tremblade, France; Cluster of Microbial Ecology, Groningen Institute for Evolutionary Life Sciences, University of Groningen, Groningen 9747 AG, The Netherlands; Department of Biology, University of Oxford, Oxford, United Kingdom; Department of Pathobiology and Population Science, Royal Veterinary College, Hatfield, United Kingdom; Department of Human Genetics, David Geffen School of Medicine, University of California Los Angeles, CA 90095, USA; Department of Biostatistics, Fielding School of Public Health, University of California Los Angeles, Los Angeles, CA 90095, USA; Department of Computational Medicine, David Geffen School of Medicine, University of California Los Angeles, CA 90095, USA

**Keywords:** landscape phylogeography, continuous phylogeography, molecular epidemiology, viruses

## Abstract

The fast rate of evolution in RNA viruses implies that their evolutionary and ecological processes occur on the same time scale. Genome sequences of these pathogens can therefore contain information about the processes that govern their transmission and dispersal. In particular, landscape phylogeographic approaches use phylogeographic reconstructions to investigate the impact of environmental factors and variables on the spatial spread of viruses. Here, we extend and improve existing approaches and develop three novel landscape phylogeographic methods that can test the impact of continuous environmental factors on the diffusion velocity of viral lineages. In order to evaluate the different methods, we also implemented two simulation frameworks to test and compare their statistical performance. The results enable us to formulate clear guidelines for the use of three complementary landscape phylogeographic approaches that have sufficient statistical power and low rates of false positives. Our open-source methods are available to the scientific community and can be used to investigate the drivers of viral spread, with potential benefits for understanding virus epidemiology and designing tailored intervention strategies.

---

Understanding how environmental factors impact the dispersal of individuals is a fundamental biological question with applications in a wide range of research fields, including ecology, conservation and epidemiology. In situations where direct observation and/or movement-tracking of individuals is not feasible, analysing the genomic variation among sampled individuals can be used to understand how environmental factors affect their dispersal, an approach which we can broadly term “landscape genetics”. First coined over twenty years ago (1), landscape genetics is the study of how landscape attributes impact the gene flow and resulting genetic variability (genetic diversity and structure) of a spatially-distributed organism (2), without requiring individuals to be assigned to discrete sub-populations. One popular population genetic approach involves analysing the association between a matrix of inter-individual genetic distances and matrices representing one or more covariates, such as environmental distances between the individuals’ sampling locations (3, 4). Environmental distances can be defined as spatial distances that are weighted according to the environmental values along a path connecting two point locations; models used to define this path include the least-cost path model (5) or models based on graph and circuit theories (3). In the context of landscape genetic analyses, these path models allow us to evaluate if environmental layers — either tested as potential conductance or resistance factors — can explain certain patterns of inter-individual genetic differentiation.

Landscape genetic analyses applied to animal and plant organisms are distinct from phylogeographic analyses that aim to study the spatial distribution and dispersal history of intra-specific (or closely related inter-specific) lineages (6, 7). Notably, when studying animal and plant species, there can be multiple generations between the occurrence of a perturbation or environmental impact and the time at which it will be detectable in the genetic variability of the studied population (8). This issue, known as the “time lag” problem, has been cited as a limitation in the field of landscape genetics (6, 8). More generally, for such organisms, landscape genetic and phylogeographic analyses differ in both temporal and spatial scales (6). For fast-evolving organisms however, this limitation does not necessarily apply. This is particularly true for numerous pathogens, such as RNA viruses, that are characterised by a rapid evolution (9), implying that evolutionary, ecological and epidemiological processes occur on commensurate time scales (10, 11). By statistically analysing the genetic differences among viruses sampled from a population, we can gain insights into the underlying processes governing their dispersal. In particular, the development of discrete (12) and continuous (13, 14) phylogeographic inference methods, as well as structured coalescent models (15, 16), have enabled the reconstruction of the dispersal history of target epidemics, and can be exploited to investigate the impact of external factors on the dispersal history and dynamic of viral lineages. This is precisely the goal of a recent field of research coined “landscape phylogeography”, which aims to explain the dynamics of phylogenetically informed-movements through external factors (17).

Landscape phylogeographic approaches are thus conceptually distinct from landscape genetics that primarily test associations between pairwise genetic and environmental distances. Furthermore, by directly analysing the association between lineage dispersal events and environmental factors, landscape phylogeographic approaches can circumvent the time lag issue explained above.

Over the last ten years, several methods have been developed to conduct landscape phylogeographic analyses. One such method is the implementation of a generalised linear model (GLM) extension (18) of the discrete diffusion model (12) in the BEAST X software package (19). This model jointly performs a discrete phylogeographic reconstruction and an analysis of covariates that are potentially associated with the transition rates between sampled locations (18). With a subsequent implementation in a structured coalescent model (20), the GLM approach has been applied in a number of studies aiming to unravel the drivers of viral epidemics of public health importance, such as the Ebola virus outbreak of 2013-2016 in West Africa (21) and the international SARS-CoV-2 circulation in Europe (22). Furthermore, several post hoc analytical procedures have also been developed to exploit the outcome of continuous phylogeographic reconstructions in order to investigate the impact of environmental factors on the dispersal location (23, 24), direction (23, 25), frequency (26, 27), and velocity (28, 29) of viral lineages.

As with most landscape genetic approaches, landscape phylogeographic analyses based on continuous phylogeographic inference (13, 14) do not require prior, arbitrary delimitation of discrete locations. This contrasts with discrete phylogeographic approaches that generally assume that all ancestors of the sampled viruses existed only at a pre-specified set of locations (17). In order to take advantage of continuous locations, a first analytical framework has been developed to assess the impact of environmental factors on the heterogeneity of the dispersal velocity of viral lineages (28, 30). In summary (see below for further details), this framework tests the correlation between inferred branch durations and their associated environmental distances (i.e. the environmental distance between branch start and end locations), which are inferred through a continuous phylogeographic analysis. While this post hoc approach can formally investigate the potential impact of environmental factors on the dispersal rate of viral lineages, it comes with potential limitations inherent to the relaxed random walk (RRW) model used to perform continuous phylogeographic inference. First, although the RRW model relaxes the assumption of a constant dispersal velocity in Brownian diffusion, it remains unknown how well this model can capture the impact of environmental factors, i.e. whether it is flexible enough to accommodate any diffusion heterogeneity imposed by environmental variables. Second, the RRW model remains restricted to dispersal processes that correlate with geographic distance, which may not always be the case, e.g., for human viruses such as influenza, whose global-scale dispersal can be better predicted by international air travel than geographic distances (18, 31).

The objective of the current study is twofold. First, we aim to overcome the two potential limitations of the *post hoc landscape phylogeographic approach* outlined above by implementing alternative *prior-informed landscape phylogeographic procedures* that integrate environmental heterogeneity before continuous phylogeographic inference is conducted. Second, we aim to implement and apply phylogeographic simulations to evaluate the statistical performance of landscape phylogeographic approaches testing the impact of environmental factors on the diffusion velocity of viral lineages, as well as the performance of another alternative approach for testing the impact of such factors on deviations from an isolation-by-distance pattern. Our analyses allow the identification of weaknesses and strengths among the different landscape phylogeographic methodologies tested, which enables us to provide clear guidelines for the scientific community interested in applying these approaches to understand the drivers of viral spread of public health and One Health importance.

## Results

### Post hoc landscape phylogeographic approaches

The first post hoc landscape phylogeographic approach evaluated in the present study is based on the **analysis of lineage diffusion velocity** and has been adapted from the previously established analytical framework introduced above (28, 30). This framework consists of a sequence of four analytical steps (28): (i) the extraction of the spatio-temporal information embedded in a series of annotated trees sampled from the posterior distribution of trees inferred through Bayesian continuous phylogeographic inference using a RRW diffusion process to model among-branch heterogeneity in diffusion velocity (13, 14); (ii) the computation of environmental distances associated with each phylogenetic branch using either a least-cost (5) or Circuitscape (3, 32) algorithm, the latter being based on circuit theory, which allows integrating a degree of uncertainty in the path taken by evolving lineages; (iii) the estimation of the correlation between the branch durations and the environmental distances computed for each phylogenetic branch; and (iv) a randomisation procedure to assess the statistical support of the correlation metric. This correlation metric is the statistic *Q*_1_ (*Q* statistic labelled “1” as a distinction to the three other *Q* statistics defined below) defined as the difference between *R*^2^_env_ and *R*^2^_null_, with *R*^2^_env_ being the coefficient of determination obtained from the linear regression between four times the branch durations *t* and the squared environmental distances *D*_env_ computed on the environmental raster (4*t* ∼ *D*^2^_env_), and *R*^2^_null_ the coefficient of determination obtained from the linear regression between four times the branch durations and the environmental distances *D*_null_ computed on the “null” raster (4*t* ∼ *D*^2^_null_). The null raster has the same dimensions as the environmental raster, but all accessible cells (i.e. land areas) have a uniform value set to “1”. Environmental distances computed from this null raster serve as a proxy for geographic distance, ensuring that the same path model — based on the least-cost or Circuitscape algorithm — is used for both environmental and geographic distances computations. As mentioned above, with these path models, an environmental raster can be tested as a potential conductance or resistance factor, i.e. increasing or decreasing the diffusion velocity of viral lineages.

The correlation statistic *Q*_1_ therefore measures to what extent taking the environmental heterogeneity into account increases the correlation between phylogenetic branch durations and environmental distances. In other words, the *Q*_1_ statistic evaluates the degree to which lineage diffusion velocity can be explained by the investigated environmental layer. In the previous version of this analytical framework (28), the statistic *Q*_1_ was defined as the difference between *R*^2^_env_ and *R*^2^_null_ where these coefficients of determination were obtained from a linear regression between branch durations and environmental distances (i.e. *t* ∼ *D*). In the updated version, however, the regression is now performed between 4*t* and *D*_2_ (4*t* ∼ *D*_2_), rather than between *t* and *D*, to align with the assessment of the diffusion coefficient identified as a metric more robust to the sampling intensity (i.e. the sampling size) than the lineage dispersal velocity (33, 34). While the lineage-dispersal velocity measures the dispersal velocity associated with a phylogenetic branch (i.e. the ratio *d*/*t* between the great-circle geographic distance *d*_gc_ travelled by a phylogenetic branch and its duration *t*), the diffusion coefficient can rather be considered as a measure of its diffusion velocity (*d*_gc2_/4*t*). An environmental factor is considered potentially explanatory only if both the associated posterior distribution of regression coefficients *β*_env_ and *Q*_1_ values are positive (29). In that case, and as described in the previous version of this analytical framework (28), the statistical support for the *Q*_1_ statistic can be formalised as a Bayes factor estimated through a randomisation procedure. This procedure involves the randomisation of phylogenetic branches within the study area while preserving the tree topology, maintaining the inferred position of the most ancestral node, and preventing nodes from falling in non-accessible (i.e. sea/water) areas (see the Materials and Methods section for further detail on this randomisation procedure).

The second post hoc landscape phylogeographic approach investigated here consists of a newly introduced **isolation-by-resistance (IBR) analysis**. Similar to diffusion coefficient metrics, isolation-by-distance (IBD) signal metrics have been found robust to the sampling effort (i.e. the sampling size) and are complementary to diffusion coefficient metrics (33): while a diffusion coefficient measures the dispersal capacity of viral lineages, an IBD signal metric measures to what extent phylogenetic branches are spatially structured, i.e. the tendency of phylogenetically closely-related tip nodes to be sampled from geographically close locations (33). The IBD signal can, for instance, be estimated using the Pearson correlation coefficient between the patristic and log-transformed great-circle geographic distances computed for each pair of tip nodes (33). Directly related to the notion of IBD signal, we propose assessing the IBR signal for a given environmental factor, i.e. the association between the phylogenetic (i.e. patristic) and log-transformed environmental distances between each pair of tip nodes. In other words, we assess how the investigated environmental factor could have impacted the IBD signal. The notion of IBR (3) is conceptually different from the notion of isolation-by-environment (IBE), which refers to a pattern where genetic differentiation increases with environmental *differences* (rather than *distances*) independently of geographical distance (35).

Similar to the analyses of lineage diffusion velocity, the IBR analyses were based on the estimation and the assessment of the statistical support of a correlation statistic *Q*. Here, this alternative *Q* statistic (*Q*_2_) was defined as the difference between (i) the coefficient of determination *R*^2^_env_ obtained from the linear regression between the patristic distances and the log-transformed environmental distances computed on the environmental raster for each pair of tip nodes, and (ii) the coefficient of determination *R*^2^_null_ obtained from the univariate linear regression between patristic distances and log-transformed environmental distances computed on the corresponding null raster (where environmental distances serve as a proxy for the geographic distance, i.e. an IBD setting). In the context of an IBR analysis, the *Q*_2_ statistic thus measures to what extent a given environmental factor could potentially explain a deviation from the IBD pattern. As with *Q*_1_, the statistical support of the *Q*_2_ statistic is evaluated with the same randomisation procedure of phylogenetic branch positions within the study area, which enables the estimation of Bayes factor support (see the Materials and Methods section for further detail on this Bayes factor computation).

Since it is not based on the analysis of phylogenetic branch positions inferred through a phylogeographic reconstruction, the IBR analysis is not necessarily a *landscape phylogeographic approach*. While a landscape phylogeographic approach aims to analyse the impact of environmental factors on lineage dispersal dynamics, the IBR analysis is indeed conceptually closer to a *landscape genetic approach* aiming to uncover the environmental factors impacting the inter-individual genetic differentiation. However, because this analysis is based on pairwise patristic distances computed on (time-scaled) phylogenetic trees, we propose to characterise it as a *landscape phylogenetic approach*, i.e. an analytical approach to investigate the impact of environmental factors on the phylogenetic distance or divergence time (in the context of time-scaled phylogenetic inference) between individuals. Also in this case, we used the same randomisation procedures as for the analyses on the lineage-diffusion velocity to assess the statistical support of the *Q*_2_ statistic. Given that those procedures are based on the randomisation of branch positions inferred on the map by a continuous phylogeographic inference, the IBR analysis implemented here still requires such a phylogeographic reconstruction, even if the outcome of the phylogeographic inference itself is not involved in the computation of the *Q*_2_ statistic.

### Prior-informed landscape phylogeographic approaches

We also introduce two new landscape phylogeographic approaches characterised as “prior-informed” because the analysis of the environmental factor is integrated before the continuous phylogeographic inference. Instead of assessing the association of environmental factors with phylogeographic estimates *a posteriori* as in the post hoc landscape phylogeographic approaches described above, environmental distances are first computed between sampling locations and then used to transform the space. The rationale behind this approach is that diffusion over distances that accommodate an environmental factor impacting the diffusion velocity of lineages should be more regular (characterised by less diffusion rate heterogeneity), or more Brownian, than over geographic distances that ignore such a factor. The idea here is thus to also perform the continuous phylogeographic analysis in a “new space” defined by these environmental distances and then compare the heterogeneity of the branch diffusion velocity as estimated by this analysis and the one based on original sampling coordinates. Concretely, we explore two possibilities (Fig. 1): transforming the space according to a **multidimensional scaling (MDS) analysis**, also known as principal coordinates analysis (36), or according to a **cartogram transformation** (37).

**Figure 1.**
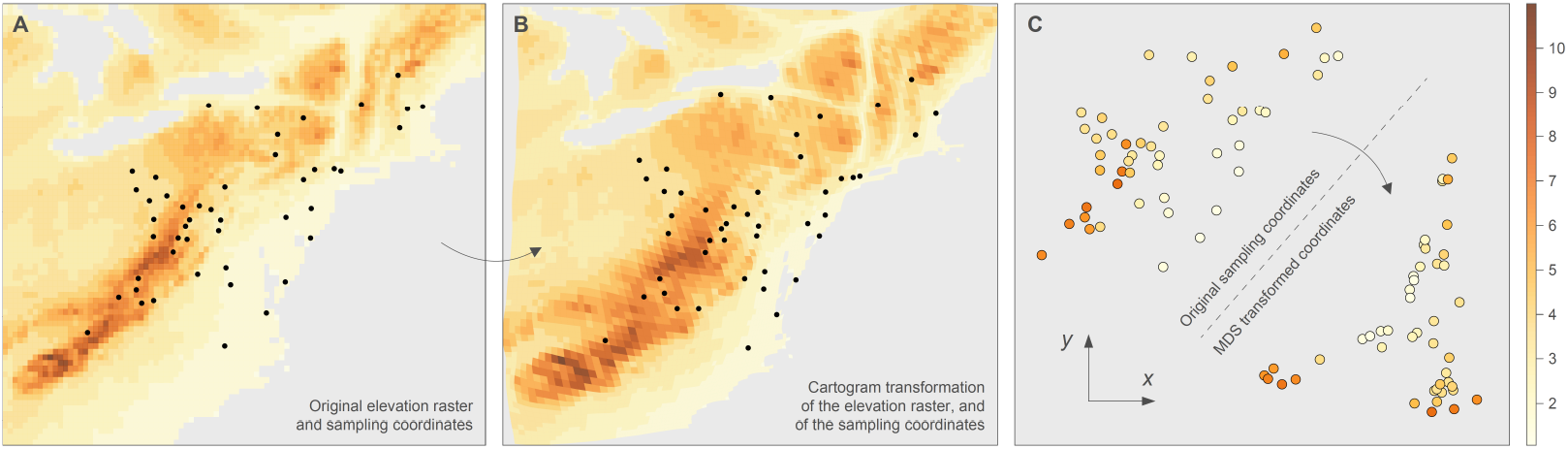
Illustration of a cartogram and multidimensional scaling (MDS) transformations based on an environmental layer tested as a potential resistance factor slowing the diffusion velocity of viral lineages. We first display the cartogram transformation (**B**) of an original elevation raster (**A**; here rescaled from 1 to 11) along with the sampling coordinates (black dots) of the raccoon rabies virus (RABV) dataset sampled in northeastern USA (38). We then also display a MDS transformation of those coordinates based on environmental distances computed with the Circuitscape algorithm among all pairs of sampling points while treating the elevation raster as a resistance factor. In this third panel, sampling points are coloured according to the altitude of their position prior to the transformation of the space. As shown in panel B, the cartogram transformation of this elevation raster — here also tested as a potential resistance factor — tends to increase the relative distance between sampling points located in higher altitude areas while decreasing the relative distance between sampling points located in lower altitude areas. As shown in panel C, a similar observation can be made when visually inspecting the relative distance among sampling points before and after the MDS transformation: sampling points in relatively low altitude areas tend to be closer to each other after the MDS transformation. The rationale behind these two prior-informed landscape phylogeographic approaches based on a transformation of the study space (with a cartogram transformation or with a multidimensional scaling analysis) is to assess if such a transformation would lead to a more Brownian diffusion process than in the untransformed study space, which would then indicate an association between the tested environmental factor and the diffusion velocity of viral lineages.

The MDS transformations are based on pairwise environmental distances computed among sampling coordinates. The sampling coordinates in the new two-dimensional MDS space (Fig. 1) are subsequently used in the continuous phylogeographic analysis. In the present study, we conduct MDS transformation based on environmental distances between sampling locations, computed using either the least-cost (5) or Circuitscape (3, 32) algorithm. As illustrated in Figure 1, the principle of cartogram transformations is to transform the size of a series of joint polygons according to values assigned to these polygons. Analogous to the MDS transformation, a cartogram transformation increases the new Euclidean distances between sampling locations associated with relatively high environmental distances, and decreases the new Euclidean distances between sampling locations associated with relatively low environmental distances. We can then assess whether the diffusion process in the transformed space, i.e. defined by environmental distances (MDS transformation) or environmental values (cartogram transformation), indeed required less heterogeneity than a diffusion process in a regular geographic space. MDS transformations have previously been applied in the context of discrete (39) or continuous (40) phylogeographic analyses of viruses although, in the latter case, they have not been used to test the impact of continuous environmental layers on the diffusion velocity of viral spread.

For the analyses based on the MDS transformations, the heterogeneity of branch-diffusion velocities was again measured through the estimation of a *Q* statistic (*Q*_3_) here defined as the difference between (i) the coefficient of determination *R*^2^_env_ obtained from the linear regression between four times the branch durations *t* and the squared Euclidean distances *d*_Eucl,env_ travelled by each phylogenetic branch in the space transformed according to environmental distances computed on the environmental raster (4*t* ∼ *d*^2^_Eucl,env_), and (ii) the coefficient of determination *R*^2^_null_ obtained from the linear regression between four times the branch durations *t* and the squared Euclidean distances *d*_Eucl,null_ travelled by each phylogenetic branch in the space transformed according to environmental distances computed on the corresponding null raster (4*t* ∼ *d*^2^_Eucl,null_). As for the analyses based on a cartogram transformation, the heterogeneity of branch-diffusion velocities was this time measured with the estimation of *Q*_4_ defined as the difference between (i) the coefficient of determination *R*^2^_env_ obtained from the linear regression between four times the branch durations *t* and the squared Euclidean distances *d*_Eucl,env_ travelled by each phylogenetic branch in the space transformed according to a cartogram transformation, and (ii) the coefficient of determination *R*^2^_null_ obtained from the linear regression between four times the branch durations *t* and the squared great-circle geographic distances *d*_gc_ travelled by each phylogeny branch in non-transformed (4*t* ∼ *d*^2^_gc_), i.e. following a standard phylogeographic reconstruction based on the non-transformed sampling coordinates. As for the *Q*_1_ and *Q*_2_ statistics, the statistical support of *Q*_3_ and *Q*_4_ is again formalised as a Bayes factor and evaluated with the procedure consisting of randomising phylogenetic branch positions within the study area.

### Simulation-based assessment of statistical performance

To evaluate the statistical performance of the four landscape phylogeographic approaches compared in this study, we applied these different methods to simulated datasets generated by two different types of continuous phylogeographic simulations involving or not the impact of an environmental factor on the lineage dispersal dynamics: (i) birth-death simulations based on a relaxed random walk (RRW) diffusion process, where the diffusion velocity is impacted by the environmental layer (hereafter referred to as “RRW simulations”), with simulation examples displayed in Figure 2 resulting in spatially-annotated time-calibrated phylogenies; and (ii) genome-sequence simulations along an actual maximum clade credibility (MCC) tree, with branch lengths modified according to various scenarios (hereafter referred to as “MCC simulations”; see the Materials and Methods section for further detail on these two simulation frameworks). The implementation of these two distinct simulation frameworks was motivated by their complementarity: while the MCC simulations involved a deterministic impact of the environmental factor, the RRW simulations involved an intrinsic stochastic impact of the environmental layer on the diffusion velocity of evolving lineages. Furthermore, in MCC simulations, genomic sequences were simulated stochastically, allowing us to investigate the impact of the Bayesian phylogenetic inference. Finally, unlike the RRW simulations, the MCC simulations included a scenario with no impact of the environmental factor on the diffusion velocity of viral lineages, which allowed testing the risk and rate of type I errors (false positives) associated with the four landscape phylogeographic approaches.

**Figure 2.**
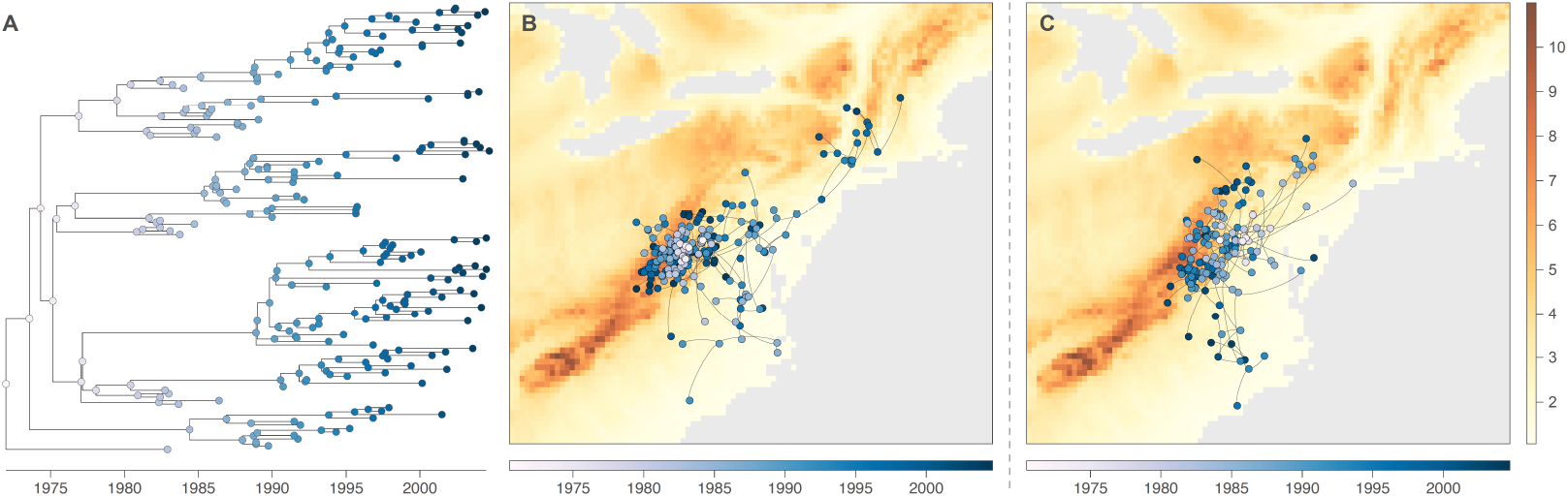
Example of continuous phylogeographic simulations based on a birth-death process and a relaxed random walk (RRW) diffusion process during which an environmental layer impacts the diffusion velocity of viral lineages. The two first panels display the phylogenetic tree obtained from a first simulation: (**A**) the time-scaled visualisation of the simulated topology and (**B**) the mapped visualisation on the environmental layer impacting the movement velocity of evolving lineages — here an elevation raster rescaled from 1 to 11 (see the Materials and Methods section for further detail) and acting as a resistance factor (i.e. slowing down movement velocity in relatively higher altitude areas). The third panel (**C**) displays the mapped visualisation of a second example of simulation. In all panels, tree nodes are coloured according to time, with internal and tip nodes coloured according to their age and collection time, respectively. The RRW simulation process illustrated here is inspired by the phylogeographic analysis of the raccoon rabies virus spread in North America (13, 30, 38) and has been used in the present study to assess the statistical performance of four landscape phylogeographic approaches aiming at testing the impact of environmental factors on the diffusion velocity of viral lineages.

For the RRW simulations, we implemented a diffusion process with simulation parameters (starting time and position, sampling time window) based on the study of rabies virus (RABV) spread in a North American raccoon population (38). The raccoon RABV dataset, published in 2007 by Biek and colleagues (38), consists of 47 sequences of ∼2,800 nucleotides (covering the complete rabies nucleoprotein gene and a large portion of the glycoprotein gene). This dataset was used to infer the dispersal history of viral lineages through a continuous phylogeographic analysis (13, 30), which subsequently highlighted the potential impact of elevation on the velocity of this viral spread (28, 30, 38). We therefore used an elevation raster acting as a resistance factor during the RRW diffusion process. The dispersal velocity of evolving lineages was negatively associated with the underlying elevation values, meaning that the lineages tended to disperse slower in areas associated with relatively higher altitudes. We conducted 50 simulations with this first RRW simulation framework. As for the MCC simulations, we again considered the raccoon RABV dataset mentioned above (38) and the actual MCC tree obtained from the continuous phylogeographic inference based on that dataset (30). These MCC simulations consisted in two successive steps: (i) modifying the branch lengths of the MCC tree according to three distinct scenarios (strong, moderate, and no impact of an underlying environmental factor on the diffusion velocity of lineages), and (ii) simulating RABV genomic sequences along the resulting time-scaled phylogeny.

The analytical workflow implemented and applied to evaluate the statistical performance of the four landscape phylogeographic approaches is summarised in Figure 3 and consists of four steps: (i) continuous phylogeographic simulations following the RRW or MCC simulation framework introduced above, (ii) MDS and cartogram transformation of the sampling coordinates for the prior-informed landscape phylogeographic analyses, (iii) continuous phylogeographic reconstructions (either based on non-transformed sampling coordinates or those transformed at the previous step), and (iv) the four landscape phylogeographic analyses.

**Figure 3.**
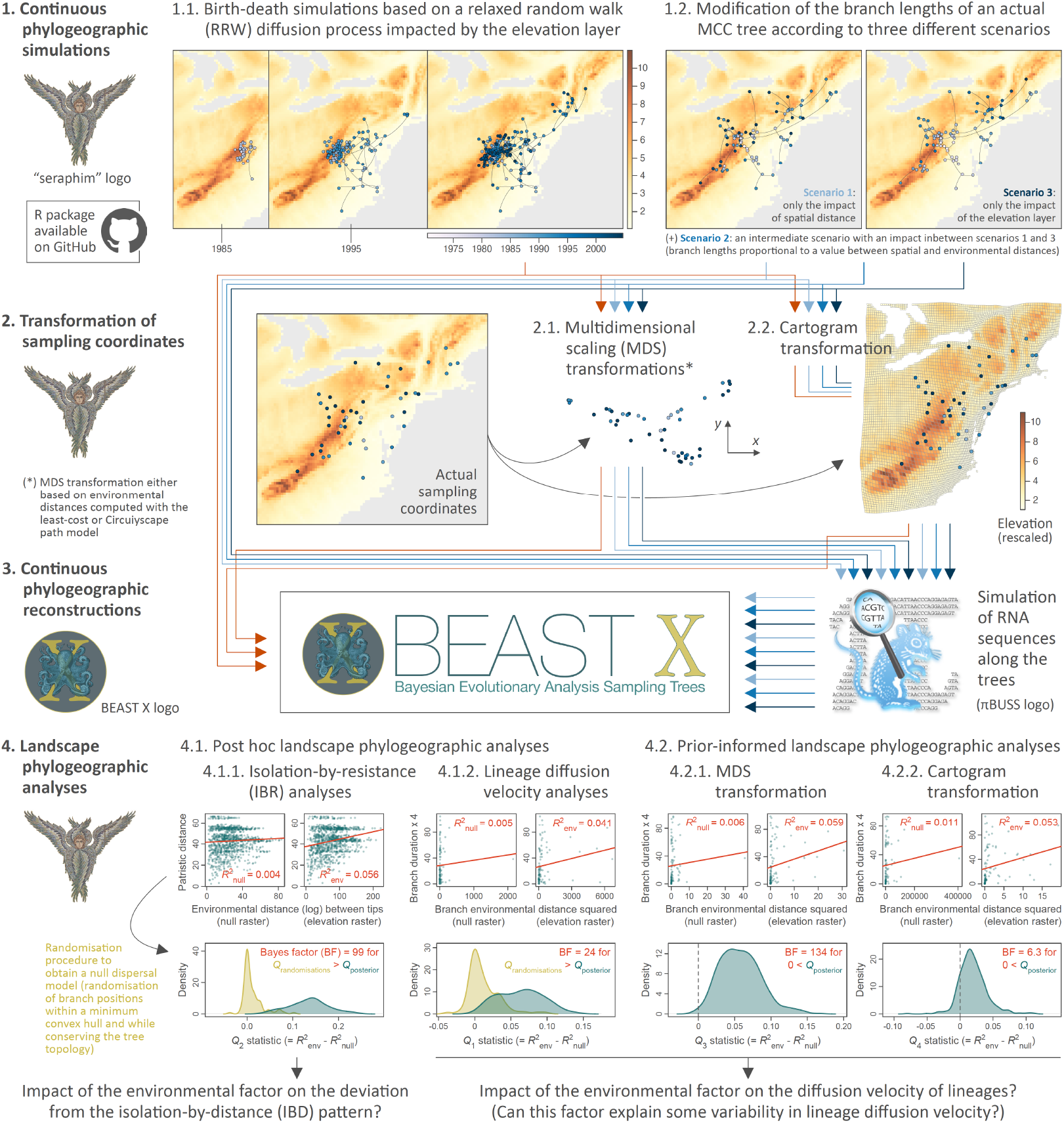
Simulation and analytical workflow implemented to assess the statistical performance of four landscape phylogeographic approaches to test the impact of environmental factors on the diffusion velocity of viral lineages or the deviation from an isolation-by-distance pattern. We here assessed the performance of two post hoc (isolation-by-resistance and lineage diffusion velocity analyses) and two prior-informed landscape phylogeographic approaches (analyses of continuous phylogeographic reconstructions based on sampling coordinates obtained either after a multidimensional scaling or cartogram transformation). Continuous phylogeographic simulations (step 1), transformations of sampling coordinates (step 2), and landscape phylogeographic analyses were conducted with the R package “seraphim” (41), the RNA sequences simulations with the program πBUSS (42), and the continuous phylogeographic reconstructions (step 3) with the software package BEAST X version 1.10.5 (19). (*) Multidimensional scaling (MDS) transformations were conducted based on environmental distances among pair of tip nodes, which were computed using either the least-cost (5) or Circuitscape (3, 32) path model while either considering the environmental raster (here an elevation raster with values rescaled between one and eleven) or a corresponding “null” raster with accessible raster cell values uniformly equal to “1” (see the text for further detail). “MCC tree” refers to a maximum clade credibility tree.

Importantly, our assessment of the statistical performance of the landscape phylogeographic approaches revealed that none seemed to be associated with detectable type I errors. When the tested environmental factor did not impact the diffusion velocity of lineages (MCC simulations under scenario 1), no type I errors were detected by any of the approaches (Table 1). Regarding the statistical power to detect a true environmental impact on lineage diffusion velocity (RRW simulations, MCC simulations with scenario 1 and, to some extent, scenario 2), our analyses led to three main results. First, the analyses revealed a notably low statistical power for the prior-informed landscape phylogeographic analyses based on a cartogram transformation (Table 1). This prior-informed approach did not show any statistical power when tested on MCC simulations conducted under scenario 2 and, most importantly, under scenario 3. Second, the RRW simulations, but also the MCC simulations with scenario 2, revealed a relatively low statistical power for the IBR approach, which we will further discuss below in light of the difference between what is actually simulated and tested by this approach. Finally, our RRW and MCC simulations indicated a moderate and good statistical power, respectively, for the two remaining landscape phylogeographic approaches, i.e. the post hoc approach based on the analyses of lineage diffusion velocity and the prior-informed approach based on a MDS transformation of sampling coordinates. However, we observed a difference when considering either the least-cost or Circuitscape algorithm used to compute the environmental distances associated with phylogenetic branches in the post hoc approach and among sampling coordinates to transform the space in the prior-informed approach. Specifically, when the Circuitscape path model is used, the statistical power tends to drop slightly for this post hoc approach when assessed under scenario 3 of the MCC simulations, and tends to notably drop for both these two post hoc and prior-informed approaches when considering the RRW simulations (Table 1). However, the MDS prior-informed approach does not seem to be impacted by the choice of the path model when assessed with the MCC simulations conducted under scenario 3.

**Table 1.**
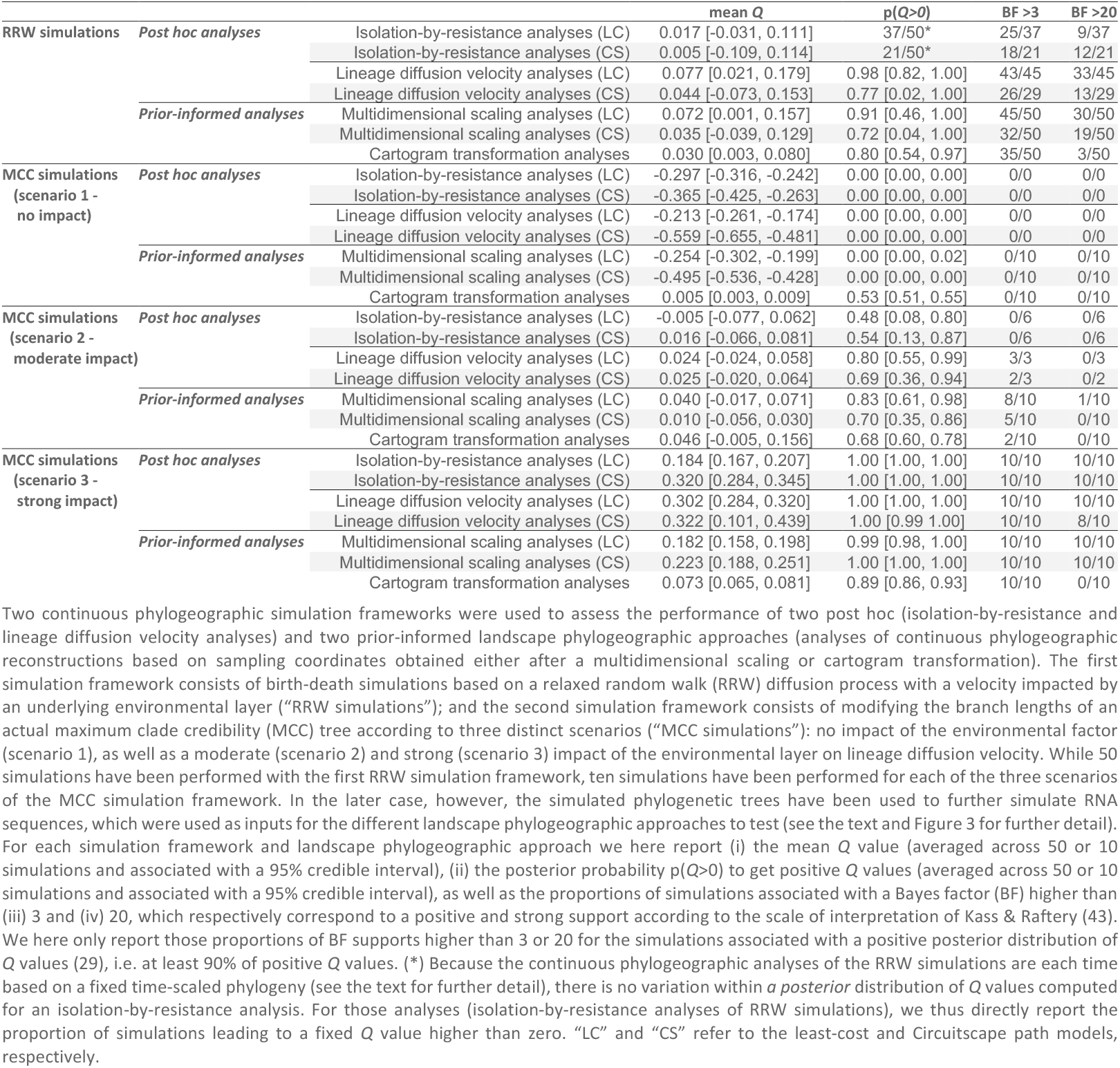
Statistical performance of landscape phylogeographic approaches for testing the impact of environmental factors on the diffusion velocity of viral lineages or on the deviation from an isolation-by-distance pattern.

## Discussion

Our assessment demonstrates that both post hoc landscape phylogeographic approaches exhibit reasonable statistical performance, with an absence of type I errors as well as good and moderate-to-low statistical power when assessed with the MCC and RRW simulations, respectively. The statistical power is particularly low for the IBR approach when evaluated with the RRW simulations. However, this result is unsurprising given the nature of these simulations, which aim to induce an impact of the environmental factor on the diffusion velocity of lineages rather than directly on the deviation from an IBD pattern, which is precisely what the IBR analyses measure. As argued earlier, the IBR analyses should be considered more as a landscape phylogenetic approach located between landscape genetic and landscape phylogeographic approaches. Previous studies (33, 34) have highlighted two types of dispersal metrics estimated by continuous phylogeographic inference, which appear to be robust to sampling intensity: the diffusion coefficient (14, 44) and the IBD signal (33, 45) metrics. While the first post hoc landscape phylogeographic approach is based on the former, the second specifically aims to investigate the cause of a deviation from an IBD pattern. Therefore both types of metric and their related landscape phylogeographic approaches can be used as they address complementary aspects of the dispersal dynamics of viral lineages.

In this study, we introduced two new landscape phylogeographic approaches based on a preliminary analysis and integration of the tested environmental factor. These approaches compare two continuous phylogeographic analyses: one based on original coordinates and one based on transformed coordinates. In the case of the MDS transformation approach, original sampling coordinates are replaced by coordinates that are transformed according to environmental distances computed on the null raster, providing a relevant proxy for pairwise geographic distances under the null dispersal model (see above). The aim is to assess whether the phylogeographic reconstruction based on the environmentally-transformed coordinates leads to more homogeneous diffusion velocities among phylogenetic branches. While both the cartogram and MDS transformation approaches do not lead to type I errors, only the analyses based on a MDS transformation reveal satisfactory statistical power (i.e. likelihood of detecting that an environmental factor does indeed affect lineage diffusion velocity). As illustrated by the cartogram transformation considered here, and reported in Figure 1, this transformation is likely not profound enough to induce a notable signature and impact of the environmental layer on the transformed sampling coordinates. Consequently, even if the environmental factor actually impacted the diffusion velocity of viral lineages, the continuous phylogeographic analysis based on those transformed coordinates appears less likely to lead to a difference in terms of homogeneity of the lineage diffusion velocity as compared to the continuous phylogeographic reconstruction based on the original sampling coordinates.

Based on our assessment of the two new prior-informed landscape phylogeographic approaches, we thus recommend the use of the MDS instead of the cartogram transformation approach. Leaving aside the IBR approach – which tests a distinct aspect of the dispersal dynamic of lineages (see above) – we provide two complementary approaches that have different benefits and weaknesses. The post hoc nature of the approach based on the analysis of lineage diffusion coefficients allows it to be applied to readily available continuous phylogeographic reconstructions that may have been previously conducted, e.g. in the context of previous studies, without any prior assumption about the impact of an environmental factor on the dispersal dynamic of inferred lineages. On the other hand, such a continuous phylogeographic analysis does not integrate the environmental heterogeneity and relies on the flexibility of the RRW diffusion model to capture the potential signature of an environmental impact on the heterogeneity of diffusion velocity among phylogenetic branches, which is not guaranteed. In light of this limitation, the prior-informed approach based on an MDS transformation presents an interesting alternative, although it comes with an important computational burden: for each environmental factor to be tested, a distinct continuous phylogeographic reconstruction has to be conducted using the transformed sampling coordinates according to this environmental factor (and see also below regarding the different rescaling parameter *k* values to test).

The landscape phylogeographic approaches evaluated here have other limitations. First, as currently implemented, they only allow testing one particular rescaling and/or transformation of the original environmental raster. In previous applications of the post hoc landscape phylogeographic approach, analysing the lineage diffusion velocity of lineages (24, 27, 46), this issue was partially circumvented by testing a series of linear transformations of the original raster cell values with the following formula: *v*_t_ = 1 + *k*(*v*_o_/*v*_max_), where *v*_o_ and *v*_t_ are the original and transformed cell values, respectively, and *v*_max_ is the maximum cell value recorded in the raster. The rescaling parameter *k* thus allows the definition of different strengths of raster cell conductance/resistance relative to the conductance/resistance of a raster cell with a minimum value set to one (24). Note that in the context of the analyses performed here, we chose to consider only a linear transformation of the original raster with a value of *k* = 10, because this corresponds to the rescaling parameter *k* value that was used to rescale and generate the environmental raster used in our phylogeographic simulations. However, in the landscape genetic literature, other approaches have been proposed, such as using a genetic algorithm to simultaneously optimise multiple environmental rasters to generate a composite resistance raster (47). Such an approach would, however, be computationally too demanding in the case of the post hoc landscape phylogeography methods considered here, as it would have to be repeated on multiple trees sampled from the posterior distribution to account for the uncertainty in Bayesian phylogeographic inference. As for the prior-informed approach based on MDS transformations, this procedure would not be relevant because it would not have any response variable to consider for its optimisation. Second, the approaches assessed here cannot analyse time-series environmental factors. This limitation is inherently associated with the algorithms used to compute environmental distances. As currently implemented, these algorithms work only on a two-dimensional grid. To enable the use of time-series environmental rasters, these algorithms would need to be adapted to work in three dimensions, allowing for the processing of successive environmental rasters over time. Finally, it is important to note that our landscape phylogeographic analyses were based on simulated datasets composed of a relatively limited number of sequences (only 47 in the case of the MCC simulations) relative to present-day genomic datasets generated in the context of viral epidemics. This choice was motivated by practical limitations related to the non-negligible computational burden of all our analyses. However, we might expect increased statistical power when these methods are applied to much larger datasets based on more comprehensive sampling.

In conclusion, our study provides guidelines on the landscape phylogeographic approaches that can be used to investigate the impact of environmental factors on the diffusion velocity of viral lineages: (i) a post hoc approach relating phylogenetic branch durations and environmental distances, updated for this study, and (ii) a prior-informed approach using MDS transformation of sampling coordinates. These two approaches are complementary given their respective conceptual and practical advantages/disadvantages, as outlined above. In addition to these two landscape phylogeographic approaches, we also introduced a third approach based on an isolation-by-resistance analysis (which might be more accurately categorised as a landscape phylogenetic approach) to investigate the environmental factors that could explain significant deviations from an isolation-by-distance pattern. Collectively, these three approaches are all available in the updated version of the R package “seraphim”, and can be applied to gain insights into the impact of environmental factors on the dispersal dynamic of fast-evolving pathogens, thereby increasing our understanding of their ecology and epidemiology.

## Materials and Methods

### Continuous phylogeographic simulations

RRW simulations were implemented in the function “simulatorRRW3”, now included in the R package “seraphim” (41); the functions “simulatorRRW1” and “simulatorRRW2” having been implemented for previous studies: the “simulatorRRW1” was designed for conducting RRW simulations along time-scaled phylogenies (26), while “simulatorRRW2” allowed birth-death simulations based on a Brownian random walk or relaxed random walk (RRW) diffusion process, without any impact of an underlying environmental factor (33). The new simulator implemented in the function “simulatorRRW3” extends the capabilities of “simulatorRRW2” by incorporating the effect of an environmental factor on the RRW diffusion process. It conducts forward-in-time joint simulations of both time-scaled phylogenies and the dispersal history of their branches on an underlying geo-referenced grid (raster) whose environmental values impact the dispersal rate of lineages. The environmental raster considered here is an elevation raster where the values had been rescaled between one and eleven using a rescaling parameter *k* value set to ten in the following formula: *v*_t_ = 1 + *k*(*v*_o_/*v*_max_), where *v*_o_ and *v*_t_ are the original and transformed cell values, respectively, and *v*_max_ is the maximum cell value recorded in the raster. At each time step of those RRW simulations, both the longitudinal and latitudinal displacements of evolving lineages are randomly drawn from a Gaussian distribution whose standard deviation is proportional (in the case of a conductance factor) or inversely proportional (in the case of a resistance factor) to the underlying raster cell value, and this while preventing lineage dispersal in inaccessible raster cells.

MCC simulations were implemented in two successive steps consisting of (i) modifying the branch lengths of an actual MCC tree according to distinct scenarios and (ii) simulating genomic sequences along the resulting time-scaled phylogenies. Branch lengths of this MCC tree were modified according to three distinct scenarios: (1) a no impact scenario, “scenario 1”, where branch lengths are proportional to environmental distances computed with the Circuitscape path model (3, 32) on a “null” raster (i.e. a raster within which the accessible cells have a uniform value set to “1”); (2) an intermediate scenario, “scenario 2”, where branch length values are randomly drawn from the branch length values generated for scenarios 1 and 3; and (3) a strong impact scenario, “scenario 3”, where branch lengths are proportional to environmental distances computed with the Circuitscape path model on an environmental raster (here an elevation raster rescaled between one and eleven). For the first and third scenario, branch lengths were set to a value proportional to the squared environmental distance — either computed on the null (scenario 1) or environmental (scenario 3) raster — divided by four, which led to a diffusion velocity dictated by the environmental distance. The program πBUSS (42) was then used to simulate genomic sequences along the resulting time-scaled phylogenies, with ten genomic datasets simulated per scenario. We simulated alignments of 12,000 nucleotides, roughly corresponding to full RABV genomes, using an HKY+Γ substitution model and an uncorrelated relaxed clock with an underlying log-normal distribution.

### Continuous phylogeographic reconstructions

All continuous phylogeographic inferences were conducted using the relaxed random walk (RRW) diffusion model (13, 14, 48) implemented in the software package BEAST X (19) version 1.10.5 (version compiled from the BEAST X GitHub repository on December 30, 2024). Specifically, for all these continuous phylogeographic analyses, we used a gamma distribution to model the among-branch heterogeneity in diffusion velocity (14). The continuous phylogeographic analyses based on the datasets obtained through RRW simulations were all based on fixed tree topologies: for each simulated dataset, we provided as a fixed tree topology the time-scaled phylogeny directly retrieved from the corresponding birth-death simulation. These phylogeographic analyses therefore did not consider sequence evolution and the program BEAST was only used to infer the ancestral position of the tree internal nodes. For the“MCC simulations”, continuous phylogeographic analyses were based on genomic sequences simulated with the program πBUSS (42) along MCC trees whose branch lengths had been modified according to the three distinct scenarios (see above). For all the continuous phylogeographic analyses based on simulated genomic sequences, we systematically modelled branch-specific evolutionary rates according to a relaxed molecular clock with an underlying log-normal distribution (49) and the nucleotide substitution process according to a GTR+Γ parameterisation (50). As for the tree prior, we specified a flexible skygrid model (51).

For each phylogeographic analysis, we used the program Tracer 1.7 (52) to assess the convergence of the Monte Carlo Markov chain (MCMC) and its mixing properties, ensuring that effective sample size (ESS) value greater than 200 for all relevant continuous parameters. We adapted the MCMC length of each analysis accordingly: all the analyses were initially run with a maximum MCMC length set to 5x10^7^ iterations for RRW simulations and 10^8^ iterations for MCC simulations. However, some analyses required re-running with a longer chain length to achieve ESS values above 200 for all relevant continuous parameters. For all the analyses, we sampled posterior trees every 10^5^ iterations and eventually discarded the first 10% of sampled trees as burn-in. We used the “postTreeExtractions” function in the R package “seraphim” (41) to extract the spatiotemporal information embedded within the 100 trees sampled from each post-burn-in posterior distribution of trees.

### Post hoc and prior-informed landscape phylogeographic analyses

The post hoc landscape phylogeographic analyses, based on measures of viral lineage diffusion velocity and isolation-by-resistance (IBR), were conducted using the method in the updated “spreadFactors” and the new “isolationByResistance” functions implemented in the R package “seraphim” (41). As for the transformations of sampling coordinates according to an MDS and a cartogram transformation, they were respectively conducted with the new functions “mdsTransformation” and “cartogramTransformation” also implemented in the R package “seraphim”. The first function calls the “cmdscale” function implemented in the R package “stats” and the second function conducts a cartogram transformation of the environmental raster (here an elevation raster) with the rubber sheet distortion algorithm of Dougenik and colleagues (37) implemented in the R package “cartogram”. All post hoc landscape phylogeographic analyses were conducted by considering either the least-cost (5) or Circuitscape (3, 32) algorithm to compute the environmental distances associated with phylogeny branches (when analysing the lineage diffusion velocity) or between each pair of tip nodes (for the IBR analyses). Similarly, the MDS transformations were based on environmental distances computed on the null or environmental raster — here an elevation raster treated as a resistance factor — either with the least-cost or Circuitscape algorithm. Overall, in this study, all the environmental distances were either computed on a null raster with a value of “1” assigned to all accessible raster cells or on an elevation raster where the values had been rescaled between one and eleven. Computing environmental distance on a rescaled environmental raster with a minimum cell value set to “1” ensured direct comparison with the environmental distances computed on the corresponding null raster, i.e. the same raster in the absence of environmental heterogeneity according to the target environmental factor.

The statistical support for the *Q* statistics estimated for the two post hoc landscape phylogeographic approaches (*Q*_1_ and *Q*_2_) was assessed using a randomisation procedure where phylogenetic branch positions are randomised within the study area. tThis randomisation step is implemented in the different functions of the R package “seraphim” and was used to conduct the four landscape phylogeographic analyses considered here. Specifically, if all trees sampled from the posterior distribution are randomised once, this randomisation procedure enables computing a Bayes factor (BF) support approximated by the posterior odds that *Q*_observed_ > *Q*_randomised_ divided by the equivalent prior odds (the prior probability for *Q*_observed_ > *Q*_randomised_ is considered to be 0.5): BF = [*p*_e_/(1-*p*_e_)]/[0.5/(1-0.5)] = *p*_e_/(1-*p*_e_), where *p*_e_ is defined as the posterior probability that *Q*_observed_ > *Q*_randomised_, i.e. the frequency at which *Q*_observed_ > *Q*_randomised_ in the samples from the posterior distribution. As previously described (28), this is analogous to computing BF support when two competing hypotheses exist, such as in the Bayesian stochastic search variable selection (BSSVS) procedure for discrete phylogeographic inference (12), where BF supports are used to assess the inclusion of rate parameters or predictors.

## Code and data availability

R scripts related to the analyses based on simulated datasets are all available, along with the associated input and output files, at https://github.com/sdellicour/landscape_phylogeography. Continuous phylogeographic simulations and landscape phylogeographic analyses were all conducted using new functions implemented in the R package “seraphim” available at https://github.com/sdellicour/seraphim, which now comes with new tutorials dedicated to (i) the phylogeographic simulations conducted on environmental rasters and (ii) the prior-informed landscape phylogeographic analyses based on an MDS or cartogram transformation, as well as (iii) an updated tutorial dedicated to the post hoc landscape phylogeographic analyses on the impact of environmental factors on the diffusion velocity of viral lineages (the latest tutorial also including the isolation-by-resistance analyses).

